# Formation of the pyruvoyl-dependent proline reductase Prd from *Clostridioides difficile* requires the maturation enzyme PrdH

**DOI:** 10.1101/2023.12.15.571967

**Authors:** Christian Behlendorf, Maurice Diwo, Meina Neumann-Schaal, Manuela Fuchs, Franziska Faber, Wulf Blankenfeldt

## Abstract

Stickland fermentation, the coupled oxidation and reduction of amino acid pairs, is a major pathway for obtaining energy in the nosocomial bacterium *Clostridioides difficile*. D-proline is the preferred substrate for the reductive path, making it not only a key component of the general metabolism but also impacting on the expression of the clostridial toxins TcdA and TcdB. D-proline reduction is catalyzed by the proline reductase Prd, which belongs to the pyruvoyl-dependent enzymes. These enzymes are translated as inactive proenzymes and require subsequent processing to install the covalently bound pyruvate. Whereas pyruvoyl formation by intramolecular serinolysis has been studied in unrelated enzymes, details about pyruvoyl generation by cysteinolysis such as in Prd are lacking. Here we show that Prd maturation requires a small dimeric protein that we have named PrdH. PrdH is co-encoded with the PrdA and PrdB subunits of Prd and also found in species producing similar reductases. By producing stable variants of PrdA and PrdB, we demonstrate that PrdH-mediated cleavage and pyruvoyl formation in the PrdA subunit require PrdB, which can be harnessed to produce active recombinant Prd for subsequent analyses. We further created PrdA- and PrdH-mutants to get insight into the interaction of the components and into the processing reaction itself. Finally, we show that deletion of *prdH* in *C. difficile* renders the corresponding mutant blind to proline, suggesting that this processing factor is essential for proline utilization. Due to the link between Stickland fermentation and pathogenesis, we suggest PrdH may be an attractive target for drug development.

**Significance Statement:** Energy conservation via Stickland fermentation was first described in the 1930s, yet information about the key enzyme of this process, Prd, is scarce, despite the fact that its central role in both metabolism and toxin production make it a promising potential drug target. Here we show how a small, previously overlooked protein that we named PrdH mediates the formation of the catalytically essential pyruvoyl-group in the active center of Prd. *In vivo* studies in *C. difficile* emphasize its critical importance in the utilization of proline. The known interplay between proline reduction and toxin production leads us to suggest PrdH as a potential drug target. Moreover, our findings open the door for further structural and functional studies with recombinantly produced active Prd.

## Introduction

The nosocomial anaerobic gut bacterium *Clostridioides difficile* is a leading cause of healthcare-associated infections and therefore a major burden for healthcare systems worldwide (1). Infection by *C. difficile* is closely linked to its unique metabolism (2), where one of the major processes of energy acquisition involves the coupled oxidation and reduction of pairs of amino acids in a process known as Stickland fermentation (3). While the oxidation of the first amino acid yields ATP via substrate-level phosphorylation, the second amino acid acts as an electron acceptor for the regeneration of NAD+. In addition to ATP generation in the oxidative arm of Stickland fermentation, new evidence suggests that the reductive section of the pathway contributes to ATP formation as well, feeding into an electron bifurcation system that ultimately adds to proton motive force. This has recently been demonstrated for the reduction of D-proline in *Clostridioides sporogenes* (4) and seems to explain the preference of D-proline as the reduced substrate (5).

Reduction of D-proline in Stickland fermentation leads to 5-aminovalerate and is catalyzed by the D-proline reductase complex Prd, which belongs to the small but ubiquitous family of pyruvoyl-dependent enzymes (6). In Prd, the pyruvoyl-containing PrdA subunit associates with a second, selenocysteine-containing protein termed PrdB, which is encoded in the same gene cluster. While PrdA and PrdB together are catalytically competent to perform D-proline reduction (Fig. 1*A*), they assemble into a large multimer of nearly 1 MDa molecular weight, probably representing a (PrdA/PrdB)_10_ stoichiometry (7).

**Figure 1.**
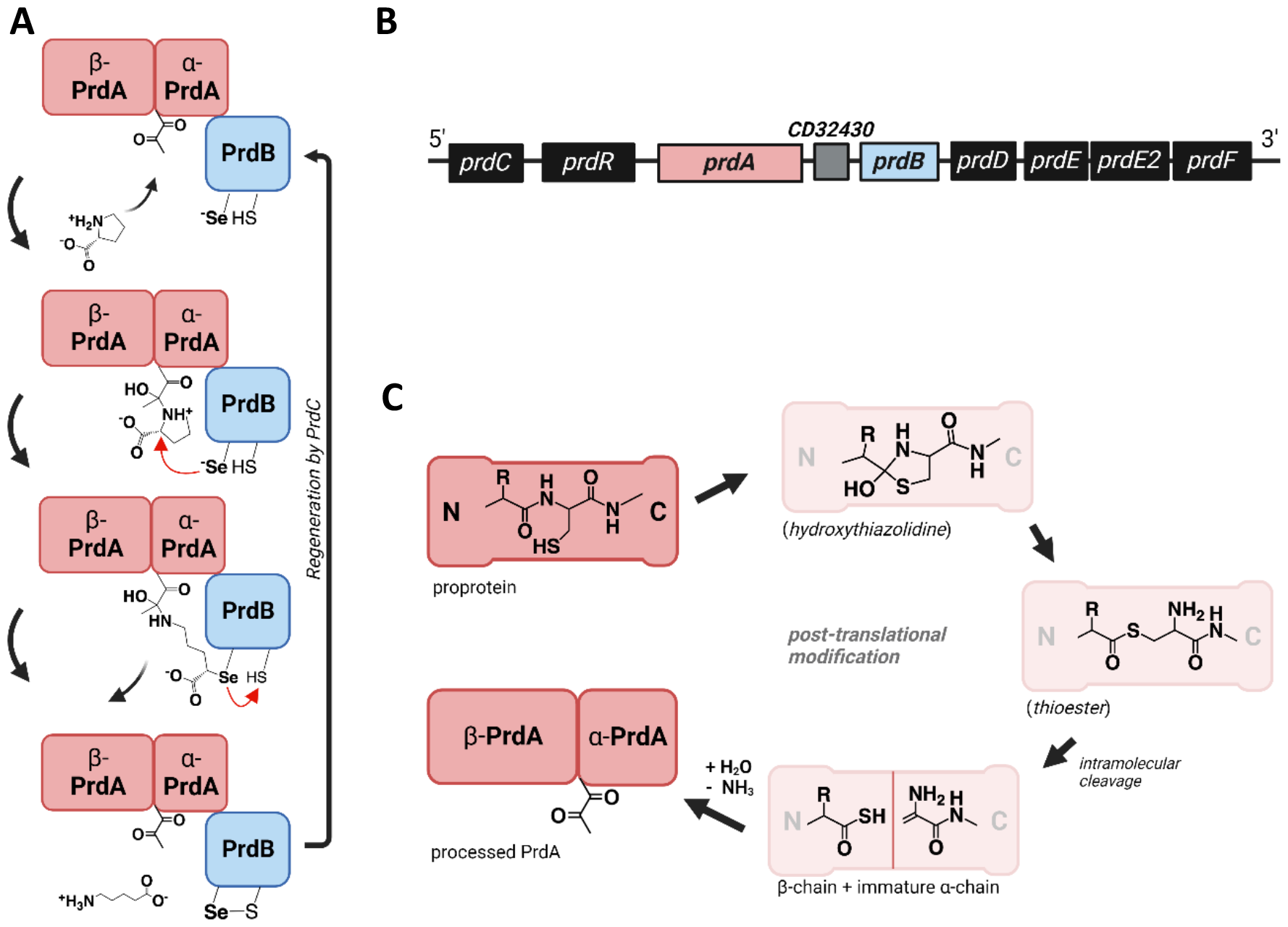
Proline reductase Prd is a pyruvoyl- and selenocysteine-containing enzyme. *(A)* Reduction of D-proline starts with binding the substrate at the pyruvoyl cofactor of theα-subunit of PrdA. The selenocysteine of PrdB attacks theα-C-atom of D-proline, which induces ring cleavage and subsequent release of the product 5-aminovalerate. Regeneration of the complex is mediated by the electron transport protein PrdC (5). *(B)* Organization of the *prd*-operon in *Clostridioides difficile*. PrdR regulates the expression of the following genes in response to the cellular D-proline level. PrdA and PrdB encode the two subunits of the proline reductase complex. PrdD and PrdE/E2 share high structural similiarities with PrdA, but their function is still unknown. PrdF encodes a proline racemase, catalyzing the conversion of L-proline to D-proline. *(C)* Proposed mechanism of pyruvoyl cofactor formation in PrdA, based on observations made for pyruvoyl-dependent decarboxylases (34).

A main characteristic of the structurally diverse pyruvoyl-containing enzymes is their translation as inactive proproteins (termed π-unit), which then mature via intramolecular cleavage at a specific sequence position with subsequent formation of a covalently bound pyruvate at the N-terminus of the C-terminal fragment, resulting in an active enzyme. Pyruvoyl-dependent enzymes can be divided into two classes. The first is represented by decarboxylases, which includes metabolic enzymes such as histidine decarboxylase, S-adenosylmethionine decarboxylase, aspartate decarboxylase and phosphatidylserine decarboxylase (6). These enzymes consist of two subunits and contain a serine that induces serinolysis at the specific cleavage site of the proprotein. The second class, to which Prd belongs, are reductases that possess a cysteine instead.

For the serine-containing pyruvoyl proenzymes, the intramolecular processing reaction is well-established (8). A related mechanism has also been proposed for the reductases, albeit that these enzymes would employ cysteine to cleave the immature protein as shown in Fig. 1*C*. Here, the thiol group of the cysteine is predicted to attack theα-C-atom of the preceding amino acid. A hydroxythiazolidine intermediate is formed and S/N-acyl shift leads to formation of a thioester intermediate. β-elimination with subsequent addition of water and loss of ammonia leaves covalently bound pyruvate at the N-terminus of the C-terminal fragment, termed theα-subunit, after cleavage from the β-subunit. Similar to the pyruvoyl-containing decarboxylases, theα- and β-subunit remain bound to each other in the reductases.

Although the mechanism of pyruvoyl formation is considered to be spontaneous, it has previously not been possible to activate recombinantly expressed PrdA under physiological conditions (9). The enzyme remained in its unprocessed state, leading to the question if an additional factor is needed for the maturation process. Recently, similar observations were made for the histidine decarboxylase (10) and the L-aspartate-α-decarboxylase (11), where such processing factors could indeed be identified.

In this study, we demonstrate that pyruvoyl maturation of the PrdA subunit of Prd is catalyzed by PrdH, a small conserved hypothetical protein co-encoded in the *prd*-operon (Fig. 1*B*). We further show that pyruvoyl formation requires the presence of PrdB, yielding stable full-length or truncated active Prd. Further experiments with mutants of PrdA and PrdH gave insights into requirements and interaction between assembly and processing reaction of the complex. In order to emphasize the crucial role of PrdH in proline reduction *in vivo*, a knock-out mutant was generated, showing that PrdH is essential for normal growth of *C. difficile* in proline-containing minimal media.

## Results

### 1. Truncation allows recombinant expression of a stable PrdA fragment in *E. coli*

The proline reductase Prd is a multimeric complex with a molecular mass of ∼960 kDa (6). Although isolation of this complex from *Clostridioides* has first been performed decades ago (12), recombinant generation of the complete complex has remained unsuccessful due to aggregation and degradation of PrdA (8). To alleviate this problem, we expressed several N-terminally truncated variants of PrdA, finding that omission of the first 148 amino acids led to a stable 52.5 kDa fragment that could be purified with chromatographic methods. Nevertheless, in line with previous studies (6), spontaneous processing to a pyruvoyl-containing protein was not observed. Instead, PrdA_149-626_ remained in its uncleaved state (Fig. 2*A*).

**Figure 2.**
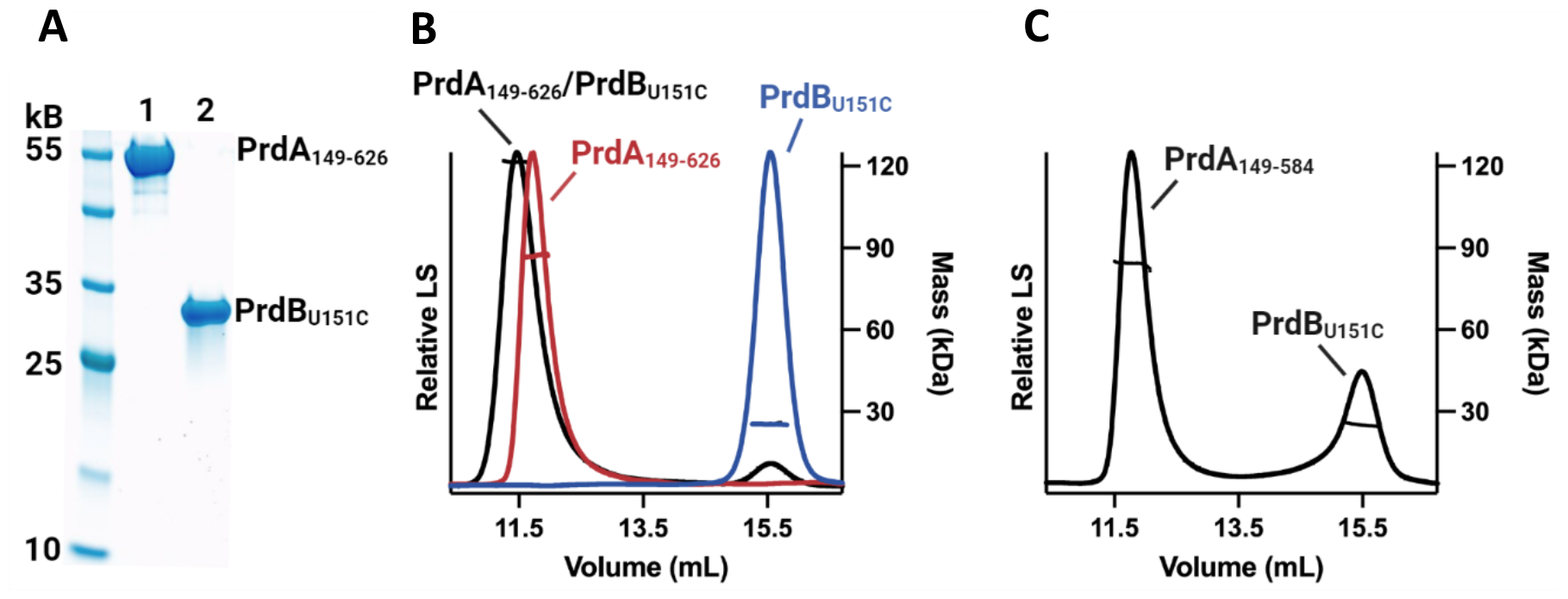
Unprocessed PrdA interacts with PrdB. *(A)* SDS-PAGE analysis of recombinant PrdA_149-626_ (1) and PrdB_U151C_ (2). *(B)* SEC-MALS results of three independently performed experiments. PrdA_149-626_ (red) elutes with a single peak. The measured mass of ∼80 kDa indicates a rapid equilibrium between monomeric and dimeric states. The measured mass of PrdB_U151C_ (∼24kDa, blue) is consistent with the calculated mass of the monomer. The mixture of PrdA_149-626_ and PrdB_U151C_ (grey) results in a shifted elution with a measured mass of ∼125 kDa, indicating strong interaction between the two subunits. Similar to PrdA_149-626_, a rapid equilibrium between different oligomeric states is assumed. *(C)* SEC-MALS analysis of a mixture of PrdA_149-584_ and PrdB_U151C_. No interaction between the two proteins is observed.

The second Prd subunit, PrdB, is a selenocysteine-containing protein, requiring the detection of a specific SECIS element in the secondary mRNA structure to recognize the UGA-codon as a signal for incorporation of selenocysteine instead of stopping the translation process (13). Despite the fact that *E. coli* produces selenocysteine-containing enzymes as well, *E. coli* did not recognize the SECIS element of clostridial mRNA in our experiments. We therefore created a PrdB variant in which we substituted the selenocysteine at position 151 with cysteine. This variant, PrdB_U151C_, has a molecular weight of 25.7 kDa and could readily be isolated using affinity chromatography (AC) followed by size exclusion chromatography (SEC) (Fig. 2A).

### 2. PrdA_149-626_ and PrdB_U151C_ interact *in vitro*

The Prd complex is composed of the two subunits of PrdA and of PrdB, suggesting that both proteins may interact tightly. To test whether the mutated and recombinantly expressed PrdA_149-626_ and PrdB_U151C_ engage in a complex despite lacking the pyruvoyl functionality in the PrdA subunit, we employed size exclusion chromatography coupled to multi-angle light scattering (SEC-MALS). The measured mass for PrdA_149-626_ alone was found to lie in between the calculated mass of a monomer and dimer, which may indicate a rapid equilibrium between both states (Fig. 2). PrdB_U151C_, on the other hand, was found to be monomeric. Mixing both subunits at a 1:1 molar ratio before SEC led to co-elution of both proteins, indeed hinting at a strong interaction. The shifted elution peak is consistent with the increased measured mass. A minor fraction of free PrdB_U151C_ was also observed. For the mixture of PrdA_149-626_ and PrdB_U151C_, the data indicate the formation of heterotetramer (PrdA_149-626_)_2_/(PrdB_U151C_)_2_ with a molecular mass of 180 kDa, as well as smaller amounts of heterotrimer (PrdA_149-626_)_2_/PrdB_U151C_ and heterodimer PrdA_149-626_/PrdB_U151C_. Larger complexes as observed for native full-length Prd isolated from *C. difficile* (see below, Fig. 7) are missing, suggesting that the N-terminus of PrdA may be crucial for higher order oligomerization.

Further, we aimed at investigating PrdA sequence motifs required for complex formation with PrdB. Toward this, we created different N-or C-terminally truncated variants of PrdA and employed SEC-MALS analysis to assess complex formation. This revealed that a relatively short 26-residue C-terminal segment of PrdA enables interaction with PrdB: while PrdA_149-612_ still binds PrdB (not shown), PrdA_149-584_ lost this ability completely despite otherwise being stable (Fig. 2*B*).

### 3. Identification of PrdH (CD32430) as a conserved protein in the *prd*-operon

The fact that recombinantly expressed PrdA remained unprocessed led us to assume that installation of the pyruvoyl moiety requires additional factors. Therefore, we analyzed the *prd*-operon and found an open reading frame downstream of *prdA*, numbered CD32430, that encodes a hypothetical protein of 94 amino acids (Fig. 1*B*).

Comparing the genome of *C. difficile* with other proline-reducing bacteria using BLAST and the STRING database (14) shows high conservation of this gene among bacteria harboring the proline reductase (Fig. S1). Available transcriptome data also show transcription of CD32430 *in vivo* (15). Due to the high conservation we decided to name the gene *prdH*.

Next, we aimed at isolating PrdH protein for further experiments. The protein was recombinantly expressed in *E. coli*, followed by purification via affinity chromatography and size exclusion chromatography. While the calculated mass of PrdH is ∼11 kDa, we observed a molecular weight of ∼22 kDa in SEC-MALS analysis (Fig. 3*A*), demonstrating that PrdH assembles into a homodimer. Because crystallization attempts where not successful, we used AlphaFold2 via ColabFold (16, 17) to construct a structural model of the dimer with a pLDDT > 90 in most regions of the protein (Fig. 3*B*). The high confidence score of this model also encouraged us to perform further structural analysis of PrdH (see section 6).

**Figure 3.**
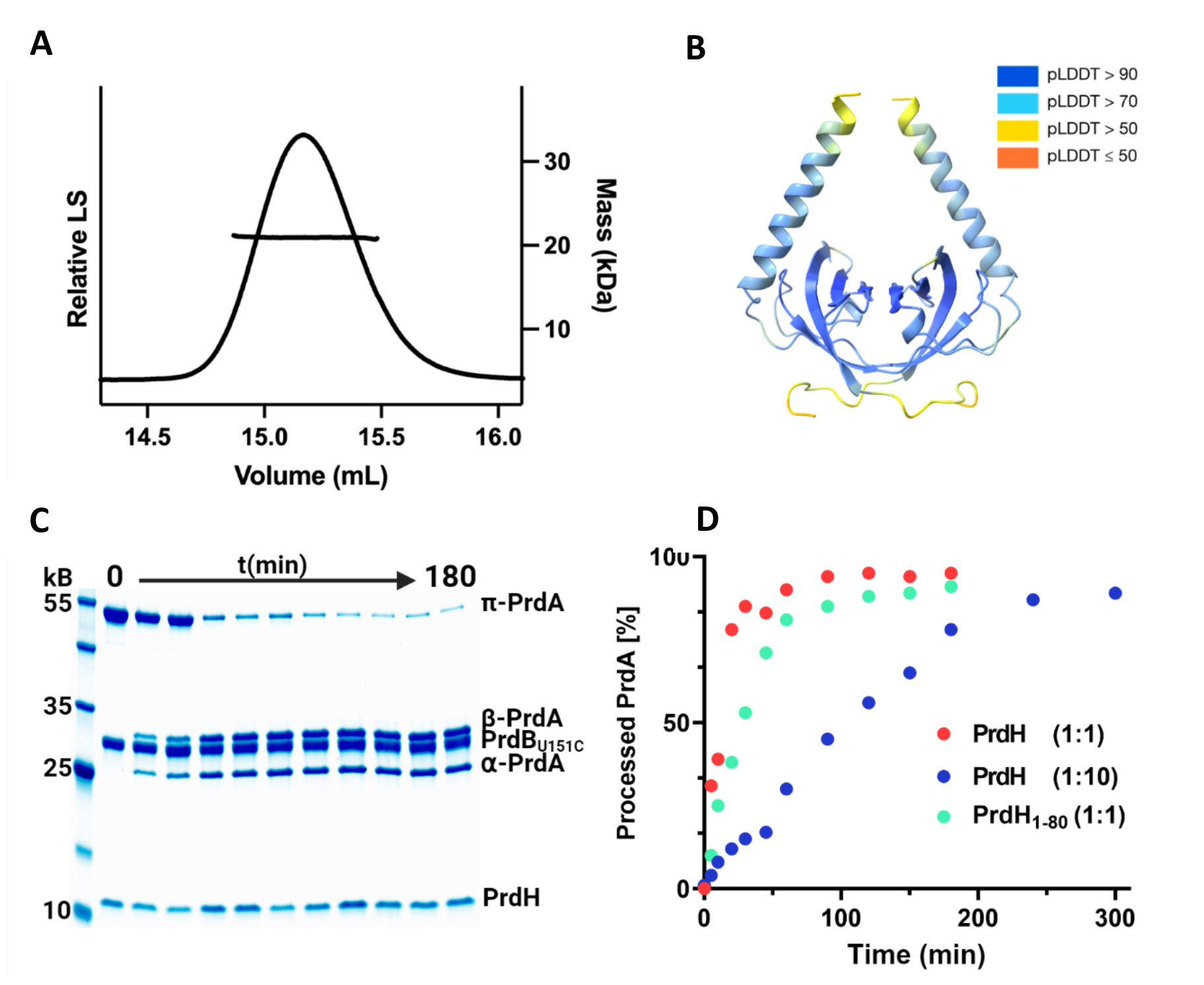
The homodimeric protein PrdH catalyzes cleavage of PrdA in the PrdA/B complex. *(A)* SEC-MALS analysis of recombinant PrdH reveals a dimeric protein. *(B)* The structure of the PrdH dimer as predicted with Alphafold2 via ColabFold (17). Colors represent the confidence score of AlphaFold2 (blue = highly confident, red = poorly confident). The image was created using ChimeraX (35). *(C)* Time-dependent *in vitro* processing of π-PrdA fragment PrdA_149-626_ in complex with PrdB by PrdH at equimolar ratio. *(D)* Band intensities were measured using ImageJ (31). The amount of processed PrdA is shown.

### 4. PrdH mediates cleavage of PrdA but requires the presence of PrdB

The maturation and concomitant installation of the pyruvoyl group in PrdA involves cleavage between T420 and C421. Although this has previously been considered to involve an autocatalytic process (18), we could not observe spontaneous cleavage in any of the recombinant fragments of PrdA (Fig. 2*A*), which is in line with previous reports (9). We therefore reasoned that installation of the pyruvoyl group may require the conserved protein PrdH as a processing factor. Incubation of PrdA_149-626_ with PrdH indeed led to cleaved protein, but to our surprise, this also required PrdB_U151C_, which needed to be included in equimolar amounts with respect to the PrdA fragment (Fig. 4*A*). Processing of PrdA appeared to require prior complex formation with PrdB, which was corroborated by the finding that PrdA_149-584_, which does not interact with PrdB (see section 2) was not cleaved in a similar experiment (Fig. 4*A*).

**Figure 4.**
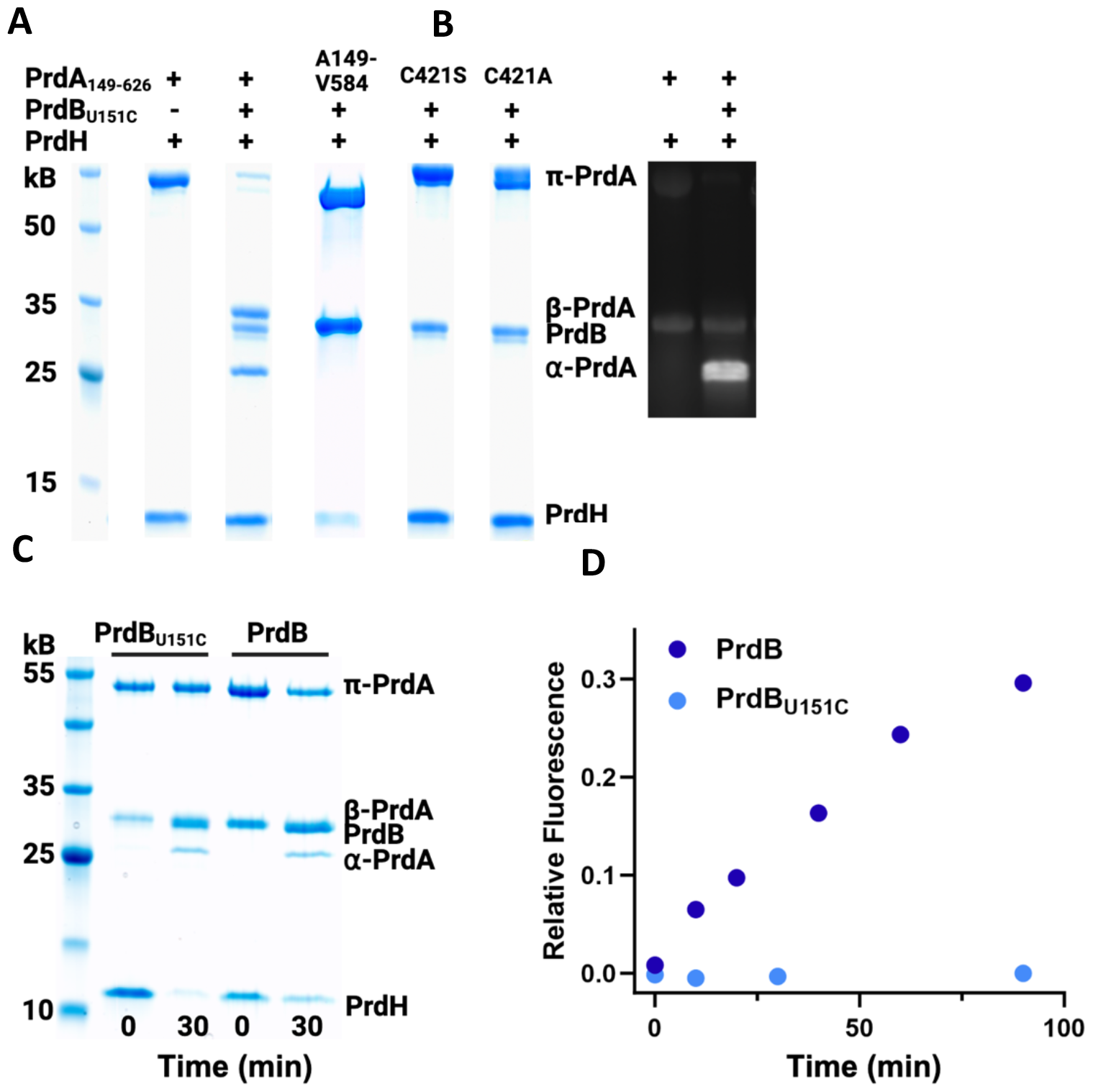
PrdH-mediated cleavage requires PrdB and C421 to install the pyruvoyl moiety in PrdA, allowing the production of catalytically active Prd fragments. *(A)* SDS-PAGE showing the requirement of a pre-formed PrdA/B complex and of PrdA-C421 for processing. Different mixtures of PrdA_149-626_ or its C421 mutants, PrdB_U151C_ and PrdH have been analyzed. *(B)* Detection of the pyruvoyl cofactor in PrdA-αvia fluorescence after incubation with fluorescein thiosemicarbazide. *(C)* Recombinant expression of selenocysteine-containing PrdB leads to the same PrdH-mediated PrdA_149-626_ processing as with PrdB_U151C_. *(D)* Detection of 5-aminovalerate via fluorescence after reaction with *o*-phthalaldehyde demonstrates the proline reductase activity of PrdA_149-626_/PrdB after processing with PrdH.

For further investigation of the cleavage reaction, a time course experiment was performed by stopping the reaction after different intervals, followed by subsequent densitometric analysis of SDS-PAGE bands. This revealed a half-life of approx. 20 minutes when PrdH was employed at equimolar concentration. Approx. 95% turnover was achieved after 2 hours. When PrdH was reduced to 10 mol-%, the half-life of PrdA_149-626_ increased to 120 minutes, but led to almost the same amount of turnover, showing that PrdH acts catalytically on the PrdA fragment (Fig. 3). Given this enzymatic activity, we next investigated if PrdH would also act on variants of PrdA in which the cysteinolysis-mediating C421 has been replaced. Both the alanine mutant PrdA_149-626___C421A_ and the serine mutant PrdA_149-626___C421S_ were not processed, indicating that cysteine is indeed required for maturation (Fig. 4*A*).

Finally, we tested if the PrdH-mediated cleavage of PrdA indeed leads to formation of the electrophilic pyruvoyl group in the C-terminal fragment. Here, we used fluorescein thiosemicarbazide, which, as a nucleophile, attacks the pyruvoyl group and leads to specific labeling of the PrdA-α-subunit as expected (19) (Fig. 4*B*).

### 5. *In vitro*-processed PrdA_149-626_ in complex with PrdB reduces D-proline

To test whether PrdA_149-626_ reduces D-proline to 5-aminovalerate after PrdH-mediated processing, *o*-phthalaldehyde, which reacts with 5-aminovalerate to form a fluorescent derivate, was used (20). Although the crucial pyruvoyl group could be clearly identified (see section 4), PrdA_149-626_/PrdB_U151C_ was not capable of catalyzing the reaction (Fig. 4*D*). We hypothesized that the selenocysteine to cysteine substitution at U151 not only reduced but also completely impeded the activity of PrdB. To circumvent difficulties of expressing selenocysteine-containing proteins in *E*.*coli* (see section 1), we used amber suppression tRNA to incorporate selenocysteine into PrdB (21). Purification of PrdB was identical to PrdB_U151C_ and incubation with PrdA_149-626_ and PrdH possessed similar processing of the PrdA fragment (Fig. 4*C*). The resulting processed PrdA_149-626_/PrdB complex indeed reduced D-proline to 5-aminovalerate (Fig. 4*D*), showing that PrdH-mediated processing of the PrdA_149-626_/PrdB complex is the only requirement to generate an enzymatically active proline reductase. Further oligomerization to the large multimeric complex found in *C. difficile*, which apparently depends on the first 148 amino acids of PrdA (see section 2), is, on the other hand, not essential for reductase activity.

### 6. Assessment of important motifs in PrdH

The fact that AlphaFold2 generates a high-confidence structural model of the PrdH dimer together with the availability of sequences from nearly 200 PrdH homologues provides an opportunity for the assessment of important sequence and structure motifs in PrdH.

First, we noted that *C. difficile* PrdH possesses a conspicuously long 25-residue α-helix at its C-terminus, which could be envisioned as a structural element that interacts with PrdA, in particular because both PrdA and PrdH are homodimeric proteins. However, shortening this α-helix by 14 residues did not affect recombinant production of the protein, and the resulting variant was not strongly impaired in catalyzing the processing of PrdA (Fig. 3D). This is in line with the finding that PrdH homologues of several other bacteria do not seem to possess this α-helix at all.

On the other hand, analysis of PrdH homologues identifies R32 as the most highly conserved residue in these proteins. Further, according to the predicted model, R32 is located at an exposed position on the surface (Fig. 5B). Indeed, mutation to alanine abrogated the activity of PrdH completely, corroborating its importance (Fig. 5C). Further experiments will be required to gain more detailed insight into the molecular mechanism underlying PrdH-mediated pyruvoyl-installation into PrdA.

**Figure 5.**
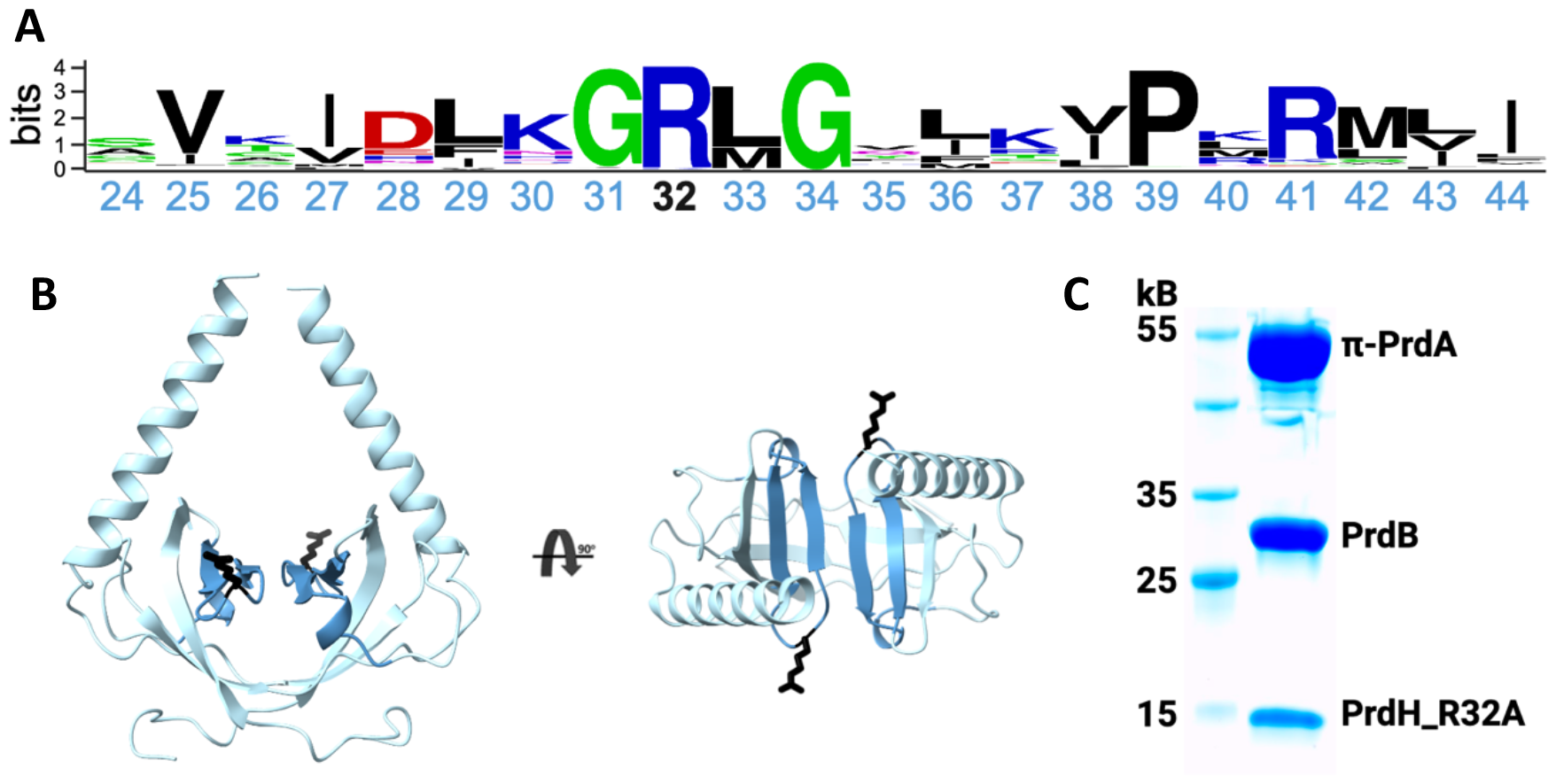
Identification of important motifs in PrdH. *(A)* Sequence conservation within a central stretch of PrdH, based on the alignment of 198 sequences (created with weblogo, (36)). *(B)* Mapping of the central stretch shown in panel A onto the AlphaFold2-model of PrdH (darker blue). The highly conserved exposed arginine R32 is colored in black. *(C)* SDS-PAGE analysis shows the requirement of R32 for PrdH-mediated processing of PrdA.

### 7. Pull-down assays reveal dynamics of the PrdA/B/H interaction

We have established that processing of PrdA requires both complex formation with PrdB and the enzymatic activity of PrdH (see section 4). Therefore, at a certain time, all three components should form a ternary complex. For further investigation, pull-down experiments using size exclusion chromatography were performed in which all three proteins were expressed separately and mixed at equimolar ratios directly before performing SEC to avoid any prior processing of π-PrdA (Fig. 6). First, PrdH co-eluted with PrdA_149-626_, indicating interaction between these two proteins, although the processing reaction requires PrdB as well (shown in section 4.) The main elution peak of a mixture of PrdA_149-626_, PrdB_U151C_ and PrdH suggests interaction between all three components, with the processing reaction occurring during the chromatography run. A mixture of PrdA_149-626_C421S_, PrdB_U151C_ and PrdH shows similar co-elution, but no processing product is observed, indicating that PrdH indeed interacts with the PrdA/B complex, but because the C421S-mutation in PrdA_149-626_ abolishes the processing reaction, its release from the PrdA/B complex is blocked. Finally, mixing PrdA_149-626_, PrdB_U151C_ and PrdH_R32A_ revealed no interaction of the PrdH-mutant with the complex at all, indicating that the highly conserved R32 plays a crucial role for the interaction.

**Figure 6.**
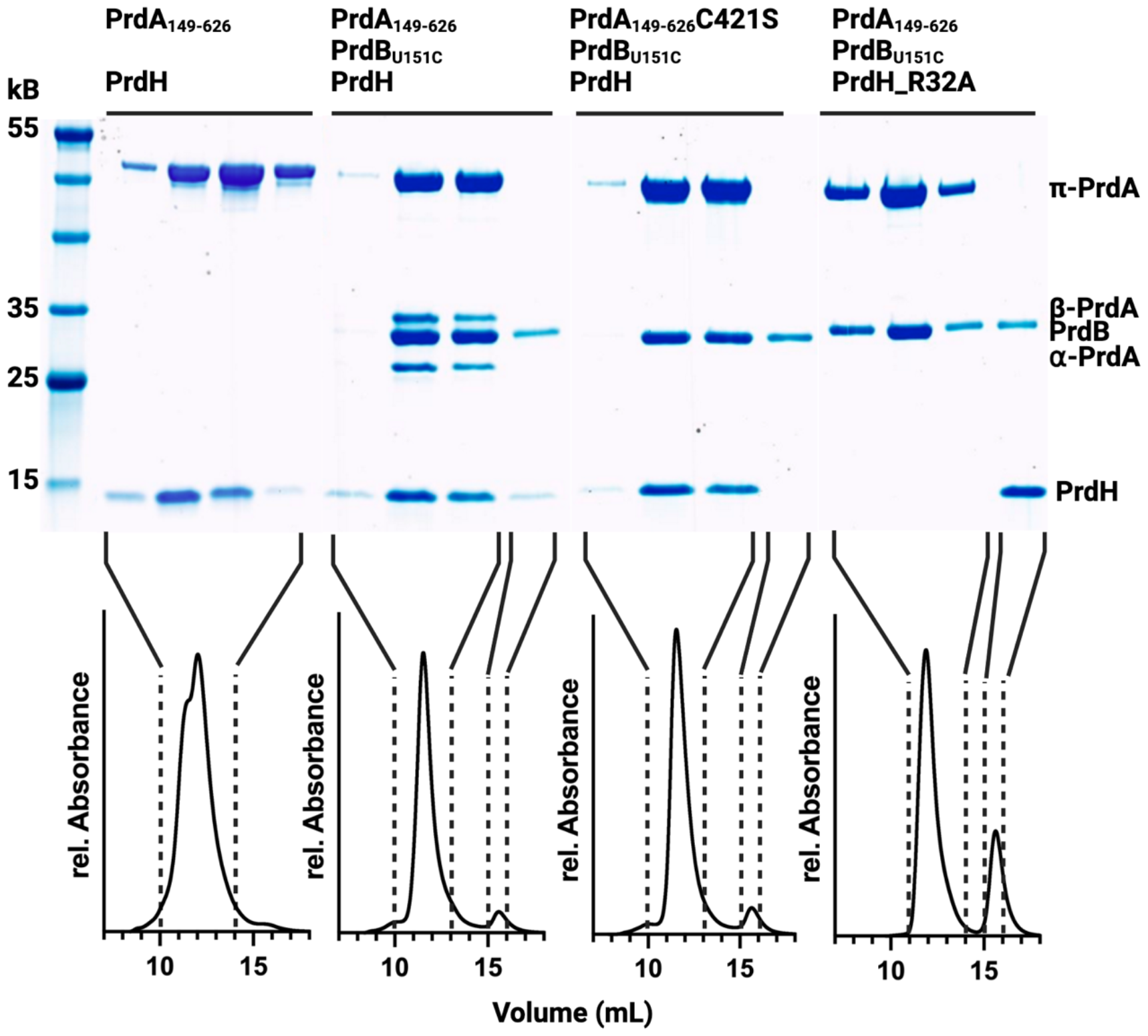
PrdA, PrdB and PrdH engage in different complexes during pyruvoyl formation. Size exclusion chromatography profiles of recombinant proteins mixed directly before analysis were recorded to identify complexes that form during the maturation of Prd. From left to right: PrdH co-eluted with PrdA_149-626_, indicating interaction between these two proteins in the absence of PrdB. Co-elution of PrdA_149-626_, PrdB_U151C_ and PrdH suggests interaction between all three components, with the processing reaction occurring during the chromatography run. While C421S-mutation in PrdA_149-626_ abolishes the processing reaction, co-elution suggests that the interaction with PrdB_U151C_ and PrdH still prevails. Mixing PrdA_149-626_, PrdB_U151C_ and PrdH_R32A_ shows that the PrdH-mutant does not interact with the complex unprocessed PrdA and PrdB at all, pointing at the critical role of R32. Note that monomeric PrdB and the homodimeric PrdH have similar molecular weight, leading to coelution without complex formation.

**Figure 7.**
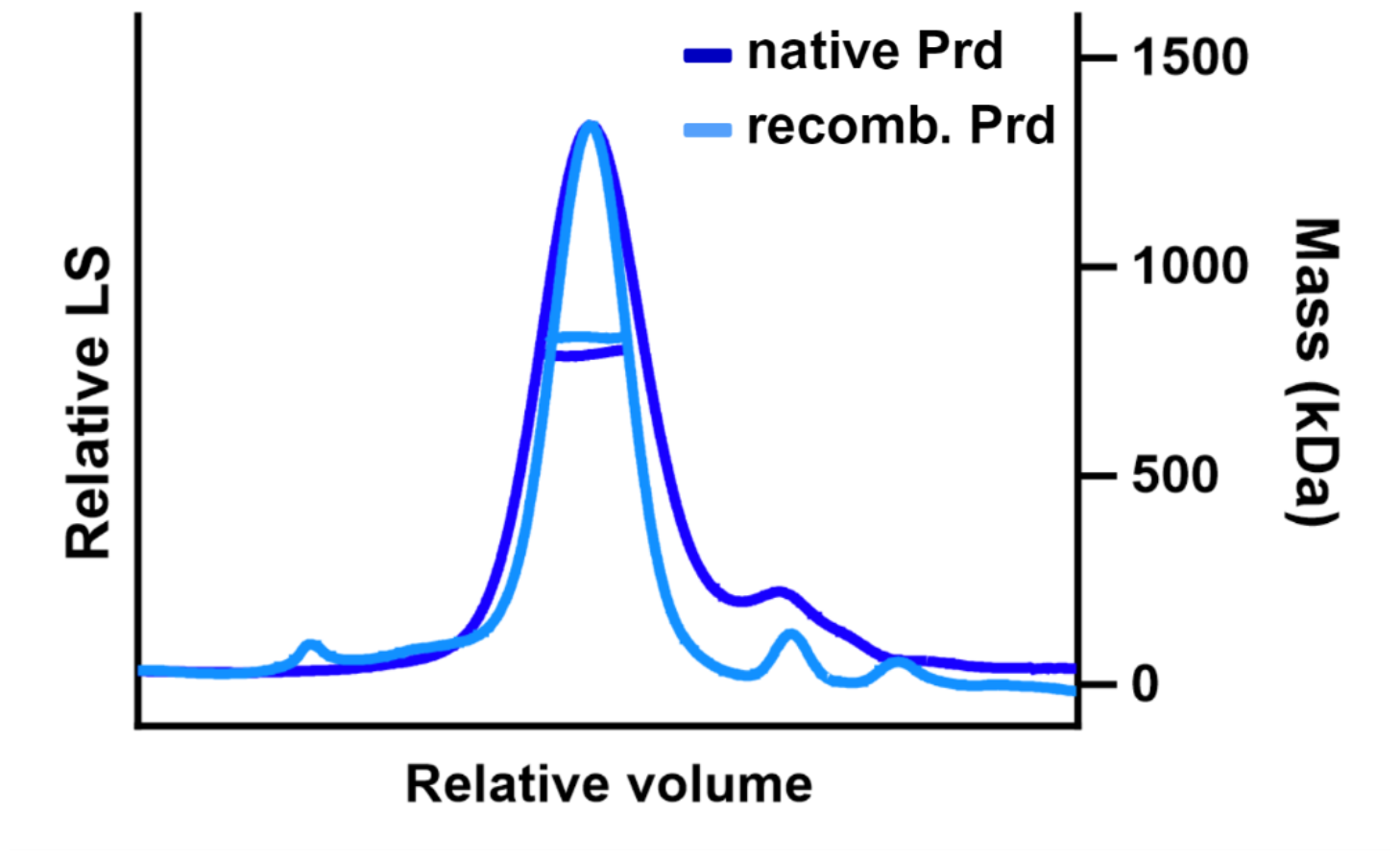
Coexpression of PrdA, PrdB and PrdH in *E. coli* leads to Prd with similar properties as Prd isolated from *C. difficile*. Comparison of SEC-MALS profiles of natively and recombinantly expressed Prd reveals similar molecular masses.

### 8. Complex assembly requires processing

The experiments summarized in the previous sections were performed with an N-terminally truncated variant of PrdA (PrdA_149-626_), which was necessary to avoid aggregation and degradation observed when full length PrdA was produced alone in *E. coli* (section 1). To our surprise, however, recombinant expression of full length PrdA became feasible through co-expression with PrdB and PrdH. Here, SEC-MALS analysis of the purified recombinant Prd reveals the same molecular mass as for native protein purified from *C. difficile*, namely approx. 960 kDa, indicating a (PrdA/PrdB)_10_ stoichiometry (Fig. 7). This not only shows that formation of the decameric Prd complex requires PrdH-mediated processing of the PrdA chain but also provides opportunities for further analysis of the complete enzyme.

### 9. The Δ*prdH* mutant of *C. difficile* is insensitive to D-proline

To determine the importance of PrdH *in vivo*, we created an in-frame deletion of *prdH* in *C. difficile* 630 (Δ*prdH*) via homologous recombination (22), to preserve expression of the remaining *prd* operon. Growth of the Δ*prdH* and the isogenic wild-type (WT) strains was assessed in *Clostridioides difficile* minimal medium (CDMM), which was supplemented with D-proline at concentrations ranging from 0 to 8 g/L (Fig. 8). In line with previous studies (5, 23, 24), *C. difficile* WT displayed proline-dependent growth, leading to higher cell density at higher proline levels. In contrast, the growth of the Δ*prdH*-mutant was found to not respond to proline. Surprisingly and against first intuition, the mutant showed similar growth rates as the wildtype strain at the highest proline concentrations, irrespective of the amount of proline present in the medium. A similar observation was made in a recent study using a PrdB-knockout (23), which suggests that PrdH is indeed essential for formation of active Prd and hence proline-dependent growth.

**Figure 8.**
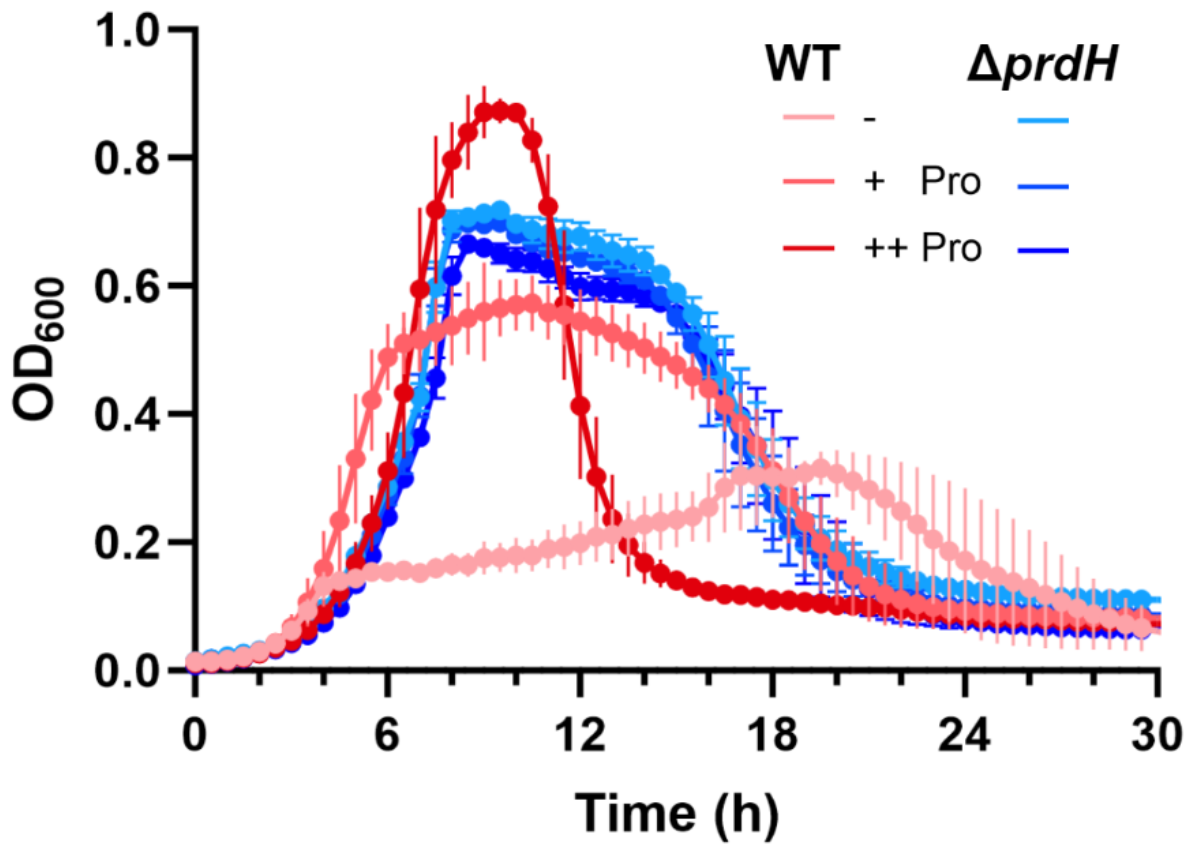
The *ΔprdH* variant of *C. difficile* is insensitive to the D-proline content of growth media. Both *C. difficile* 630 (WT) and *C. difficile* (Δ*prdH*) were grown in CDMM medium under anaerobic conditions supplemented with different concentrations of D-Proline (+ Pro: 2 g/L ; ++ Pro: 8 g/L). Data points represent the means of quadruplicate cultures with error bars representing standard deviations.

Since the utilization of D-proline for energy metabolism is connected to the production of CDI inducing toxins, with higher levels of toxins being produced at low proline concentrations (25), these findings suggest that PrdH could be a targeted to reduce the virulence of *C. difficile*.

## Discussion

Proline reductase Prd is a key enzyme of Stickland fermentation, a process that utilizes the coupled oxidation and reduction of amino acids for energy generation in bacteria such as *Clostridioides difficile*. Prd is a large multimeric enzyme consisting of PrdA and PrdB chains that contain an essential selenocysteine in PrdB and require the posttranslational installation of a pyruvoyl group via internal cysteinolysis in the PrdA subunits. While pyruvoyl moieties are also found in other unrelated proteins, details of pyruvoyl maturation in PrdA are not understood.

In this study, we show that the conserved open reading frame CD32430, renamed here as *prdH*, encodes for a homodimeric 92-residue-containing protein with crucial function for pyruvoyl formation in Prd. Using recombinant proteins, we further demonstrate that PrdA needs to undergo complex formation with PrdB before PrdH-mediated processing can occur. These findings allow the drawing of conclusions about the maturation process of the native proline reductase complex: after the translation of PrdA as an inactive proprotein, it has to interact with the separately translated PrdB first before PrdH can induce self-mediated cleavage of PrdA with concomitant pyruvoyl group formation. In this regard, the requirement of PrdB binding could be a mechanism to prevent the maturation of free PrdA, since the pyruvoyl group of such a protein would be expected to be highly reactive towards nucleophiles, leading to unspecific and potentially dangerous reactions before active D-proline-specific Prd is formed. On the other hand, our experiments with soluble fragments of PrdA show that higher order oligomerization to the 960 kDa complex, which is apparently mediated through the first 148 amino acids of PrdA, is not required to generate processed Prd subunits. Through processing of PrdA_149-626_ and incorporation of Sec into PrdB, we could further show that oligomerization is not essential for proline reductase activity, since the non-oligomerized processed protein consisting of a (PrdA_149-626_)_2_PrdB_2_ heterotetramer was fully functional in reducing D-proline to 5-aminovalerate.

Further, our findings indicate that PrdH catalyzes the processing reaction, i.e. PrdH is an enzyme. However, since the pyruvoyl installation is an intramolecular reaction conceptually not requiring catalysis by an external factor, it is at present not clear how PrdH stimulates this reaction mechanistically. Future studies will reveal if complex formation induces strain in the amino acid sequence undergoing cysteinolysis in PrdA or if PrdH participates in the reaction by e.g. acting as an acid/base catalysist. However, our finding that PrdH already interacts with PrdA in the absence of PrdB, albeit that PrdB is nonetheless required for pyruvoyl formation, suggests that PrdH and PrdB may act together in increasing conformational strain in PrdA. Such conformational strain seems to be a general requirement to induce acyl shift reactions in proteins (26), a common feature shared by diverse autoprocessing mechanisms such as protein splicing or cleavage of hedgehog protein precursors (18).

Interestingly, the PrdA_149-626_C421S_ mutant shows no processing at all, while complex formation with PrdB and PrdH was unaffected. In contrast, processed PrdA_149-626_ shows no interaction with PrdH. PrdH only interacts with unprocessed PrdA and detaches from the complex as soon as the reaction is complete. This is a prerequisite for PrdH’s ability to act as an enzyme.

The overall findings lead us to assume that PrdH is an essential enzyme for proline reduction in *C. difficile* and therefore also a potential target against CDI. The observed insensitivity of the constructed PrdH-knockout-mutant to D-proline in the growth medium confirmed this hypothesis. Further, it could be shown that the processing reaction is a prerequisite for complex assembly, allowing the stable isolation of recombinantly expressed full length Prd. This opens opportunities for further structural and functional studies of Prd, which are currently pursued in our group.

## Materials and Methods

### Bacterial strains and growth media

Genes were amplified from genomic DNA of *Clostridioides difficile* Δ630 (DSM 27543). For purification of native Prd, *C. difficile* 630Δ*erm* (DSM 28645) was used. Both strains were obtained from the DSMZ culture collection. Cloning and expression work was performed in *E. coli*. For cloning, either NEB® Turbo Competent *E. coli* (High Efficiency) or Top10 (Invitrogen) were used. For expression, *E. coli* BL21 (DE3) or One Shot™ BL21 Star™ (DE3) were used. Overnight cultures were grown in LB-medium. For protein expression, ZYM-5052 autoinduction medium (27) was used. Depending on the plasmid, either 100 mg/L ampicillin or 30 mg/L (100 mg/L for expression) kanamycin were added. *C. difficile* cultures were grown anaerobically inside a Coy chamber (85% N_2_, 10% H_2_, 5% CO_2_) in Brain Heart Infusion (BHI) broth or on BHI agar plates (1.5% agar). When required, 15 mg/L thiamphenicol, 250 mg/L cycloserine or 8 mg/L cefoxitin were added. For growth in defined *C. difficile* minimal medium (CDMM), medium was prepared as described by Neumann-Schaal et al. (28) with minor alterations: NaH_2_PO_4_ was added to a final concentration of 16.7 mM and HNaO_3_Se to a final concentration of 9.9 µM. Subcultures of four replicates were used for inoculation of main cultures with CDMM, supplemented with different concentrations of L-proline. Growth was measured using a Synergy™ H1 microplate reader.

### Plasmids

All plasmids used in this study are listed in Table S1. Cloning for recombinant expression was performed via the SLIC method (29) by employing the ClonExpress Ultra One Step Cloning Kit from Vazyme (Cat#C115).

### Protein expression and purification

Plasmid-transformed *E. coli* were grown in LB-medium at 37 °C and 130 rpm overnight. The overnight culture was used to inoculate 1 L ZYM-5052 media to an OD_600_ of 0.1. The cells initially were grown at 37 °C and 130 rpm for 3 hours and then for another 25 hours at 20 °C before harvesting by centrifugation and storing at -80 °C until further preparation.

For protein purification, the cells were resuspended in 50 mM HEPES, 300 mM NaCl, 2.5% glycerol, pH 7.5 with protease inhibitors (cOmplete mini EDTA-free, Sigma Aldrich) followed by lysis trough sonication and subsequent centrifugation at 100.000 g for 45 min. The supernatant was then applied to either Ni^2+^-charged HisTrap HP 5 mL (Cytiva) or StrepTrap HP 5 mL (Cytiva) via an ÄKTA™ pure or ÄKTA™ go system (Cytiva). Elution of bound protein was performed using elution buffer (20 mM HEPES, 300 mM NaCl, 2.5% glycerol, pH 7.5 and either 1 mM desthiobiotin for StrepII-tagged protein or 250 mM imidazole for His-tagged protein). Fractions containing pure protein as indicated by SDS-PAGE were concentrated by ultrafiltration and applied to a HiLoad 16/600 Superdex 200 (for PrdA and PrdB) or HiLoad 16/600 Superdex 75 (for CD32430/PrdH) (both Cytiva) equilibrated in SEC buffer (20 mM HEPES, 300 mM NaCl, 2.5% glycerol, pH 7.5). Fractions containing the desired pure protein as indicated by SDS-PAGE were pooled, concentrated, frozen in liquid nitrogen and stored at -80 °C for further analysis.

Native Prd was isolated by modification of a previously published protocol (30). *C. difficile* 630Δerm (DSM 28645) was cultivated anaerobically in proline-containing CDMM (0.76 g/L consisting of 0.1 g/L added to the 0.66 g/L of the casamino acids as characterized for this casamino acids lot, Merck) and harvested by centrifugation in the late exponential phase. The pellet of 1.6 L cell culture was resuspended in 40 ml lysis buffer (50 mM HEPES pH 8, 500 mM NaCl, 10 mM imidazole, 5 mM BME, DNase I, cOmplete™ EDTA-free protease inhibitor cocktail) and cells were lysed by sonication before slow addition of 40 ml ice-cold 2 M (NH_4_)_2_SO_4_. Cell debris and precipitated proteins were separated by centrifugation at 100.000xg for 1 h. Prd was then purified from the supernatant by hydrophobic interaction chromatography (5 mL HiTrap Butyl HP, Cytiva), followed by an anion exchange step (8 mL MonoQ, Cytiva) and finally size exclusion chromatography (Superose 6 10/30 increase column, Cytiva).

### Size exclusion chromatography – multi angle light scattering (SEC-MALS)

Superdex 200 Increase 10/300 or Superdex 75 Increase 10/300 (both Cytiva) columns were used with an Agilent 1260 Infinity II HPLC system coupled to a miniDAWN TREOS MALS detector and an Optilab T-rEX refractometer (Wyatt Technology Corp.). Degassed SEC buffer (20 mM HEPES, 300 mM NaCl, 2.5% glycerol, pH 7.5) was used for equilibration and 100 µg protein were applied.

### Assay of *in vitro* cleavage of PrdA

Recombinantly expressed and purified variants of PrdA and PrdB were mixed in small PCR tubes before PrdH was added. All components were present at equimolar concentration. The mixture was incubated at 37 °C for up to 3 hours. The reaction was stopped by addition of SDS loading buffer followed by heating at 95 °C for 5 minutes before applying samples to an SDS gel. For further analysis of the reaction kinetics, band intensities were measured with ImageJ (31).

### Detection of pyruvoyl cofactor

For detection of the pyruvoyl group, fluorescein thiosemicarbazide was used (19). 5 µL of protein solution (1 mg/mL) were incubated with 10 µL of 300 mM potassium acetate pH 4.6 and 1 µL of 0.1% fluorescein thiosemicarbazide in dimethylsulfoxide for 2 hours at room temperature in the dark. Separation of the proteins as well as separation of unbound fluorophore was performed via SDS-PAGE. The N-terminal carbonyl group was identified by detecting the fluorescence on a phosphorimager (Fujifilm FLA-9000).

### Detection of D-proline reduction

For detection of D-proline reduction, a fluorometric assay with *o*-phthalaldehyde as described by Seto (20) was performed. The mixture for D-proline reduction included 170 µM processed protein in 50 mM HEPES pH 7.5, 150 mM NaCl, supplemented with 10 mM MgCl_2_, 20 mM DTT and 10 mM D-proline. The reaction was stopped adding *o*-phthalaldehyde mixture, which reacts with 5-aminovalerate, building a highly fluorescent derivative. Fluorescence was measured using 340 nm for excitation and 455 nm for emission via Spark® Multimode Microplate Reader (Tecan Group Ltd.).

### Generation of a CD630_32430 deletion strain

Allelic exchange cassettes were designed with approximately 1.2 kb of homology to the chromosomal sequence flanking the deletion sites of *CD630_32430*. To avoid affecting transcription and translation of genes located upstream, the deleted region was restricted to 36 nt upstream of the CDS and 177 nt of the CDS.

Gene deletions were constructed via homologous recombination (22) and transformed into *E*.*coli* CA434 (32) with subsequent conjugation in *C*.*difficile* according to Kirk et al. (33).

## Acknowledgments

CB and MD were supported by the German Research Foundation (DFG GRK 2223: Protein Complex Assembly (PROCOMPAS)).

## Notes

**Competing Interest Statement:** The authors declare no conflict of interest.

### Competing Interest Statement

The authors have declared no competing interest.

