## Supplemental figures and tables for "Formation of the pyruvoyl-dependent proline reductase Prd from *Clostridioides difficile* requires the maturation enzyme PrdH"

**This PDF file includes:**

Figures S1 to S2
Table S1
SI References

**Fig. S1.** STRING network (1) visualizing putative interactions (edges) between proteins (nodes) based on genome wide co-occurrence. Only interactions with a high confidence score cutoff of 0.7 (default cutoff is 0.4) are shown. *(A)* PrdA from *Clostridioides difficile* 630 as input. *(B)* PrdB from *Clostridioides difficile* 630 as input. CAJ70140.1 is an alternative identifier for CD32430 (PrdH).

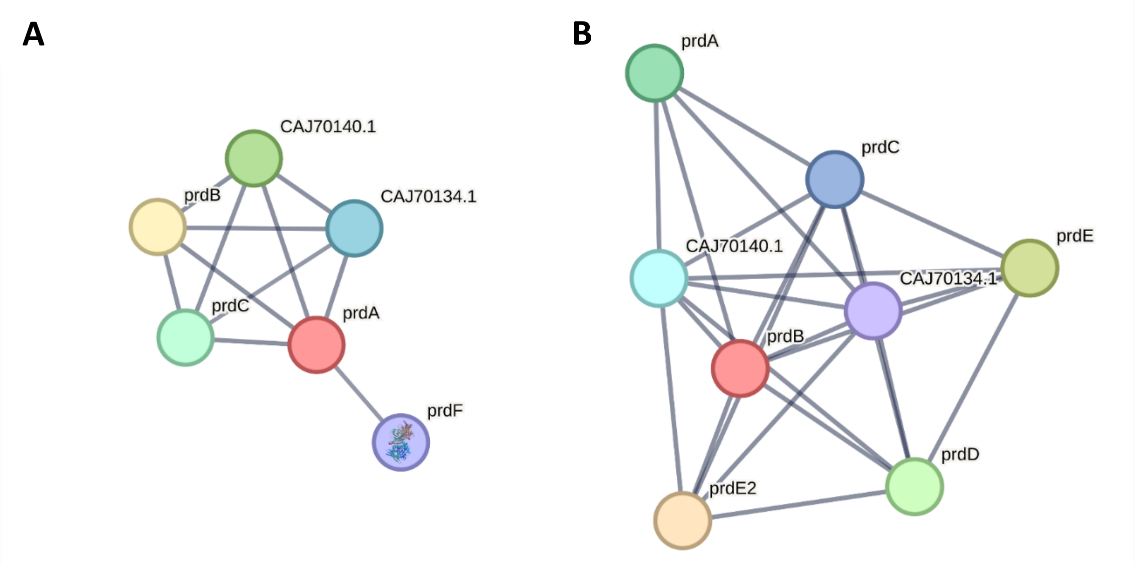

**Fig. S2.** Schematic workflow of the purification of native Prd. *(A)* 1 M $(NH_4)_2SO_4$ was added to lysed cells for precipitation, followed by centrifugation. The cleared lysate was initially purified via hydrophobic interaction chromatography (HIC) (5 mL HiTrap Butyl HP, Cytiva). The $(NH_4)_2SO_4$ concentration was reduced from 1 M to 0 M. Eluting material was analyzed by SDS-PAGE and pooled as indicated by the black box. *(B)* A finetuned KCl gradient was applied in the intermediate ion exchange step (IEX) (8 mL MonoQ, Cytiva). Different species of Prd-containing fractions were obtained and analyzed by SDS-PAGE, pooled and kept for further purification. *(C)* Prd was finally polished by size exclusion chromatography (SEC) (Superose 6 10/30 increase, Cytiva). SDS-PAGE shows three bands, corresponding to PrdA-β, PrdB and PrdA-α.

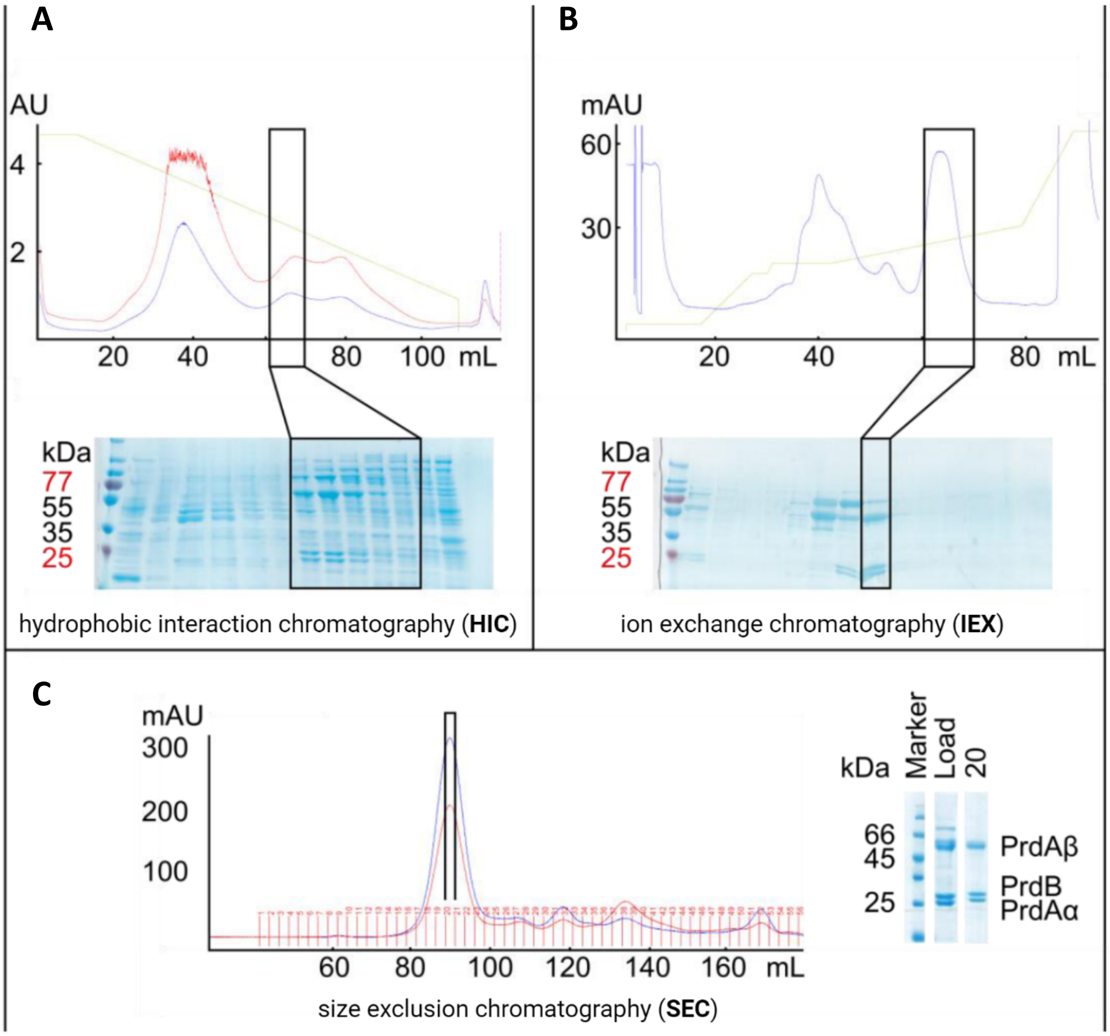

**Table S1.** Plasmids used in this study.

| Plasmid | Description | Source |
|---|---|---|
| pCB001 | pET containing $PrdA_{149-626}$ with N-terminal 6xHis-tag | This work |
| pCB002 | pET containing $PrdB_{U151C}$ with C-terminal 6xHis-tag | This work |
| pCB011 | pET containing $PrdA_{149-584}$ with N-terminal 6xHis-tag | This work |
| pCB012 | pET containing $PrdA_{149-612}$ with N-terminal 6xHis-tag | This work |
| pCB016 | pET containing PrdH with N-terminal 6xHis-tag | This work |
| pCB028 | pET containing $PrdA_{149-626\_C421S}$ with N-terminal 6xHis-tag | This work |
| pCB029 | pET containing $PrdA_{149-626\_C421A}$ with N-terminal 6xHis-tag | This work |
| pCB038 | pET containing $PrdH_{R32A}$ with N-terminal 6xHis-tag | This work |
| pCB040 | pET containing $PrdH_{1-80}$ (14 aa less) with N-terminal 6xHis-tag | This work |
| pSecUAG-Evol2 | Plasmid required for incorporation of Sec at TAG sites. | Purchased from addgene (#163148) |
| pCB043 | pET containing PrdB with TAG-codon at position 151 with C-terminal 6xHis-tag | This work |
| pFF-418 | Derived from pJAK112 (36) for generating a CD630_32430 knock-out strain | This work |
